# Majority of the highly variable NLRs in maize share genomic location and contain additional target-binding domains

**DOI:** 10.1101/2022.10.05.510735

**Authors:** Daniil M. Prigozhin, Chandler A. Sutherland, Sanjay Rangavajjhala, Ksenia V. Krasileva

**Affiliations:** Berkeley Center for Structural Biology, Molecular Biophysics and Integrated Bioimaging Division, Lawrence Berkeley National Laboratory, Berkeley, CA 94720, USA; Department of Plant and Microbial Biology, University of California, Berkeley, CA 94720, USA

## Abstract

Nucleotide-binding, Leucine Rich Repeat proteins (NLRs) are a major class of immune receptors in plants. NLRs include both conserved and rapidly evolving members, however their evolutionary trajectory in crops remains understudied. Availability of crop pan-genomes enables analysis of the recent events in the evolution of this highly complex gene family within domesticated species. Here, we investigated the NLR complement of 26 nested association mapping (NAM) founder lines of maize. We found that maize has just four main subfamilies containing rapidly evolving highly variable NLR (hvNLR) receptors. Curiously, three of these phylogenetically distinct hvNLR lineages are located in adjacent clusters on chromosome 10. Members of the same hvNLR clade show variable expression and methylation across lines and tissues, consistent with their rapid evolution. By combining sequence diversity analysis and AlphaFold2 computational structure prediction we predicted ligand binding sites in the hvNLRs. We also observed novel insertion domains in the LRR regions of two hvNLR subfamilies that likely contribute to target recogniton. To make this analysis accessible, we created NLRCladeFinder, a Google Colaboratory notebook, that accepts any newly identified NLR sequence, places it in the evolutionary context of the maize pan-NLRome, and provides an updated clade alignment, phylogenetic tree, and sequence diversity information for the gene of interest.

## INTRODUCTION

Plants rely on repertoires of innate immune receptors for protection against rapidly evolving pathogens. Nucleotide-binding, Leucine-Rich Receptor proteins (NLRs) constitute the majority of known disease resistance genes in plants (Tamborski and Krasileva 2020). Plant NLR proteins are distinguished by their central Nucleotide-Binding (NB-ARC) domains, responsible for multimerization of the receptor during activation. The specificity of NLRs is typically encoded in the Leucine-Rich Repeat (LRR) domains but can also be encoded in additional Integrated Domains (IDs) (Sarris et al. 2016; Mukhi et al. 2021; Cesari 2018). NLR signaling is dependent on NB-ARC multimerization and usually is achieved by N-terminal RPW8, Coiled Coil (CC), or TIR domains (Song et al. 2021). Recent structures of NLRs in inactive monomer (Wang et al. 2019b) and signaling-competent multimeric conformations (Wang et al. 2019a; Ma et al. 2020; Martin et al. 2020; Zhao et al. 2022; Förderer et al. 2022) have revealed the structural basis for NLR activation and signaling. They also revealed a novel non-LRR C-terminal domain present in some TIR domain-containing NLRs that contributed to target binding (Martin et al. 2020; Ma et al. 2020).

Individual NLRs differ in their intraspecies allelic diversity: some are conserved and universally present, while others show only presence-absence variation. The most sequence diverse receptors often show more complicated copy number variation along with an abundance of point mutations and small insertions/deletions, leading to a wealth of divergent alleles within the pan-genome (Baggs et al. 2017; Barragan and Weigel 2021). We have previously investigated allelic diversity in NLR immune receptors of *Arabidopsis thaliana* and *Brachypodium distachyon*, leading to the identification of highly variable NLR clades (hvNLRs) characterized by high sequence diversity on the amino acid level (Prigozhin and Krasileva 2021).

Some NLRs show high steady state expression to monitor for pathogen effectors, while others are inducibly expressed upon pathogen detection (Bieri et al. 2004; Mohr et al. 2010; Lai and Eulgem 2018). Regulation of gene expression can evolve rapidly within a species (Penning et al. 2019; Sun et al. 2023), and in the case of NLRs allow for plasticity in response to pathogen pressure through differentiation of transcript abundance, regulation of induction, and tissue specificity (Borrelli et al. 2018; Contreras et al. 2023). Tissue specific expression may be related to the pathogen specificity of NLRs, as exemplified by root-specific expression of the NRC6 cluster involved in cyst and root-knot nematode resistance in tomato (Lai and Eulgem 2018; Lüdke et al. 2023). In *Arabidopsis* Col-0, hvNLRs are more expressed and less methylated across tissues than their non-hvNLR paralogs, indicating a potential relationship between genomic features and speed of amino acid evolution at the intraspecies level (Sutherland et al. 2024).

To test if species used in agriculture have retained a similar level of NLR sequence diversity, we used protein sequences from 26 nested association mapping (NAM) founder lines (Hufford et al. 2021) to characterize their NLRs and identify the hvNLR subset in maize. We found that Rp1 (Collins et al. 1999; Ramakrishna et al. 2002), RppM (Wang et al. 2022), and RppC (Deng et al. 2022) represent three out of just four phylogenetically distinct hvNLR clades present in maize, and that these genes lie within a 2-Mb region of maize chromosome 10. Members of the same hvNLR clades show silent, tissue-specific, and constitutive expression across lines, and vary in their gene body methylation. We also combined our sequence diversity analysis with AlphaFold2 structure prediction to identify likely target binding sites in maize hvNLRs and observed presence of additional insertion domains in the maize hvNLRs. Collectively, our results show a relative reduction in NLR sequence diversity in maize that mimics the overall reduced NLR numbers in this important species coupled with genomic compartmentalization of the NLR diversity.

## RESULTS

Analysis of NB-ARC containing proteins in maize showed that NLRs are distributed over all ten maize chromosomes (Figure 1A). In order to determine which of them might belong to the hvNLR clades, we built a phylogeny of NLRs found in NAM lines based on the NB-ARC domain and rooted this tree on a branch connecting RPW8 domain-containing and CC domain-containing NLRs (Figure 1B). Initial NLR clades were selected based on bootstrap support and target clade size. The final parameters used were 14 minimum and 250 maximum clade sizes. Best supported clades were picked for every leaf, and these clades were made non-redundant by excluding smaller nested clades, leading to 88 initial clades. No bootstrap cutoff was used, instead, we tested several combinations of minimal and maximal allowed clade sizes to avoid poorly supported clades clustering together. There were 5 poorly supported clades (18-67 bootstrap support, red stars in Figure 1B), each of them could be broken down into smaller well supported clades, where at least one subclade was smaller than the 14 leaves minimum size. All of these were split into separate subclades in the following refinement step.

**Figure 1.**
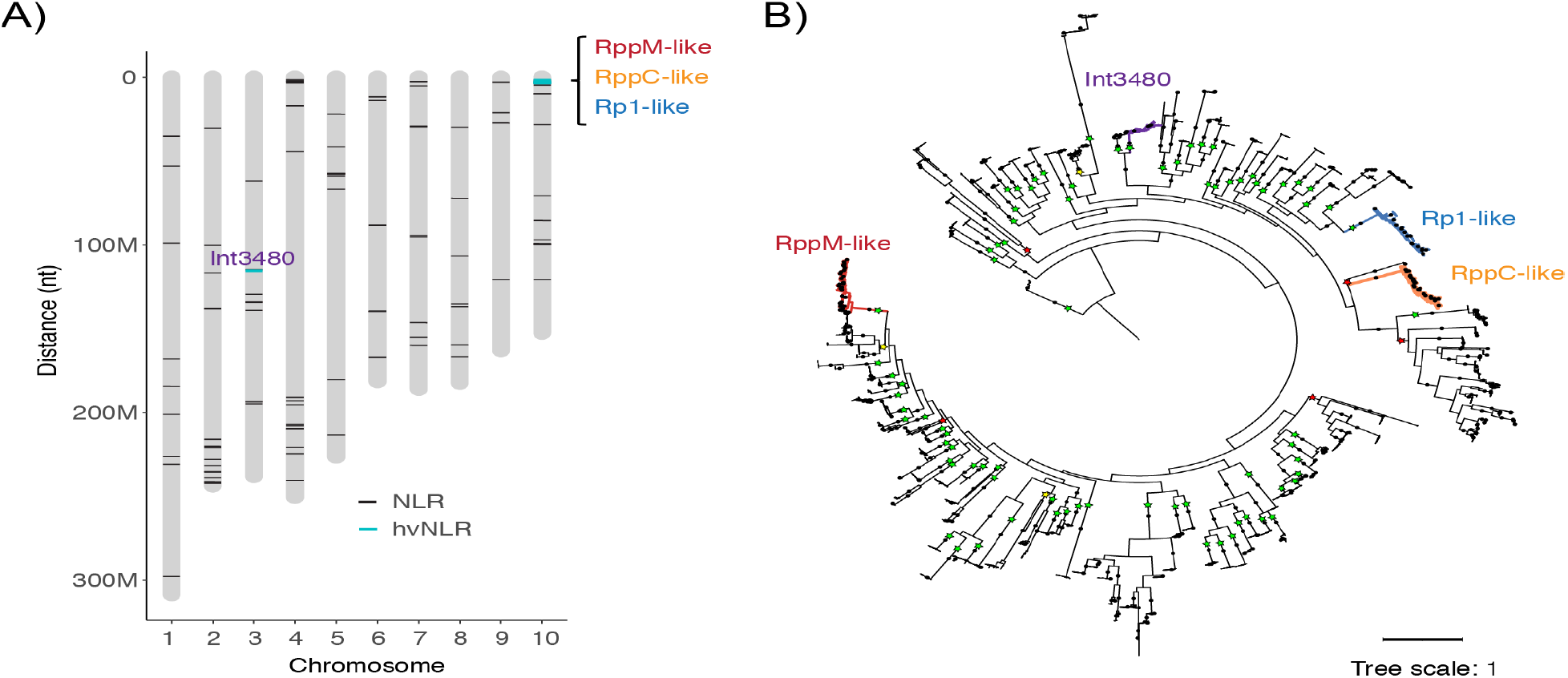
Chromosomal locations and phylogeny of maize NLR immune receptors. A) Chromosome diagram of B73 reference line showing the position of all NLR genes (black) and hvNLRs (blue) highlights a region on chromosome 10 containing an hvNLR cluster. B) Maximum likelihood phylogeny of the NLR disease resistance protein family across maize NAM founder lines. The dots on the branches signify bootstrap support greater than 90%. Initial clades that contain highly variable sub-clades are shown with colored branches. Nodes that define initial clades are labeled with stars (colored by bootstrap support: green for >90%, yellow intermediate, and red for <70%). Tree scale is average substitutions per site.

We refined the initial clades using our published pipeline based on full-length protein sequence alignments and phylogenies, then selecting long, well supported (bootstrap >90%) internal nodes to further split each clade into subclades, a process that converged after 3 iterations and resulted in 190 final clades that approximated allelic series (Supplemental Figure 1A). For each of the final clade alignments, we computed entropy scores and defined hvNLR clades as before (at least 10 amino acid sites over 1.5 bit entropy score) (Prigozhin and Krasileva 2021), which resulted in 10 candidate hvNLR clades among NAM lines of maize (Supplemental Figure 1B). After visual inspection of the resulting entropy plots, we excluded four candidate clades with signal concentrated in the very C-terminus of the alignment that did not resemble the periodicity of the signature of ligand binding sites in the LRR region (Supplemental Figure 1C). Remaining six hvNLR final clades corresponded to just four initial clades, a much lower number in comparison to 14 initial hvNLR clades in *Arabidopsis* and 18 in *Brachypodium*. Three of the hvNLR clades contained known maize resistance genes Rp1, RppC, and RppM. Our overall NLR phylogeny suggested that the four hvNLR clades are distantly related to each other (Figure 1B) and are interspersed with non-hvNLRs, similar to what we observed for hvNLR clades of *Arabidopsis* and *Brachypodium* previously (Prigozhin and Krasileva 2021).

Rp1, RppC, and RppM-like hvNLR clades clustered on chromosome 10 (Zm-B73-REFERENCE-NAM-5.0 chr10:1561295..3404163), while the remaining hvNLR clade, Int3480, was situated on chromosome 3 (Figure 1A). The roughly 2 Mb stretch of chr10s contained 4 phylogenetically distinct NLR clades including Int3985, a low variability NLR phylogenetically unrelated to any of the hvNLRs, and three hvNLR clades (Figure 2). Interestingly, we observed that the three hvNLR clades occupied distinct territories within this region, i.e. there was no Rp1-like sequence sandwiched between RppM sequences. Additionally, while all three hvNLR clades showed extensive copy number variation, members of the conserved Int3985 clade were present in the same genomic neighborhood as a single copy gene.

**Figure 2.**
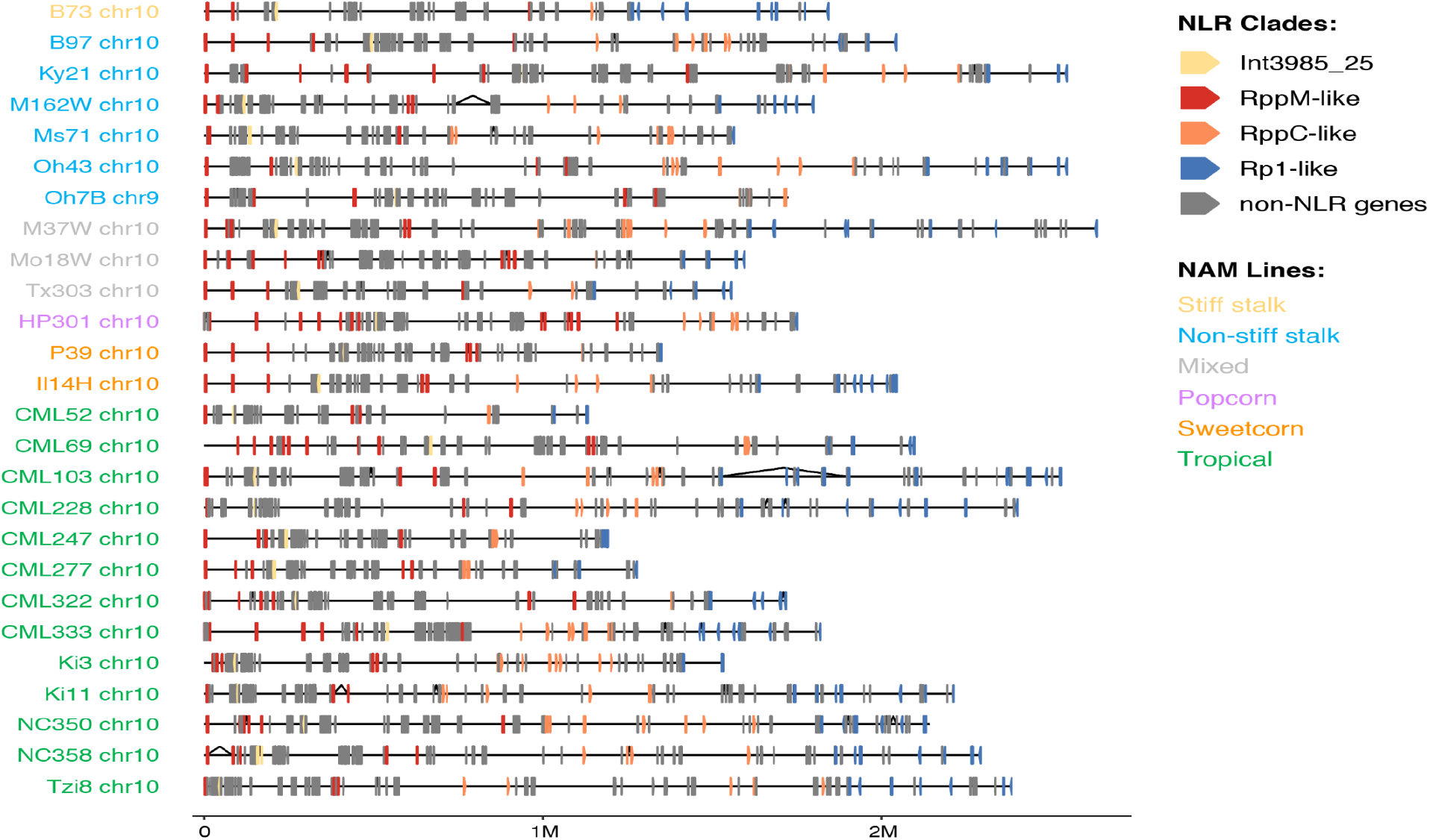
Three hvNLR clusters, along with a non-hvNLR clade, are co-located on chromosome 10. Gene diagram of hvNLR cluster in 26 NAM lines shows the ∼2 Mb region on chromosome 10 containing three of the four phylogenetically distinct hvNLR subfamilies in addition to a non-hvNLR clade, Int3985. In line Oh7B the cluster is located on chromosome 9 due to a known reciprocal translocation (Hufford et al. 2021). Black triangles denote large introns.

To investigate the expression of NLRs across the pan-genome, we used the associated RNA-seq data collected from the root, shoot, anther, ear, and the base, middle, and tip of the tenth leaf (Hufford et al. 2021). Endosperm and embryo tissues were excluded from analysis because they were not available for all NAM lines. In these pathogen unchallenged tissues, 11% of all maize NLRs are expressed constitutively, 43% show tissue-specific expression, and 42% are silent (Figure 3A). We observed within-clade variation in NLR expression, with members of the same clade showing different patterns of expression across lines. This was true for all four hvNLR clades (Figure 3A; 3B). Of the hvNLR clades, RppC-like had the highest median expression, followed by RppM, Int3840, and Rp1 (Figure 3B). RppC and RppM-like clade members show high steady state expression and are in the top 25% of all NLR expression in leaf tissue in 24/26 and 18/26 NAM founder lines, respectively (Supplemental Figure 2).

**Figure 3:**
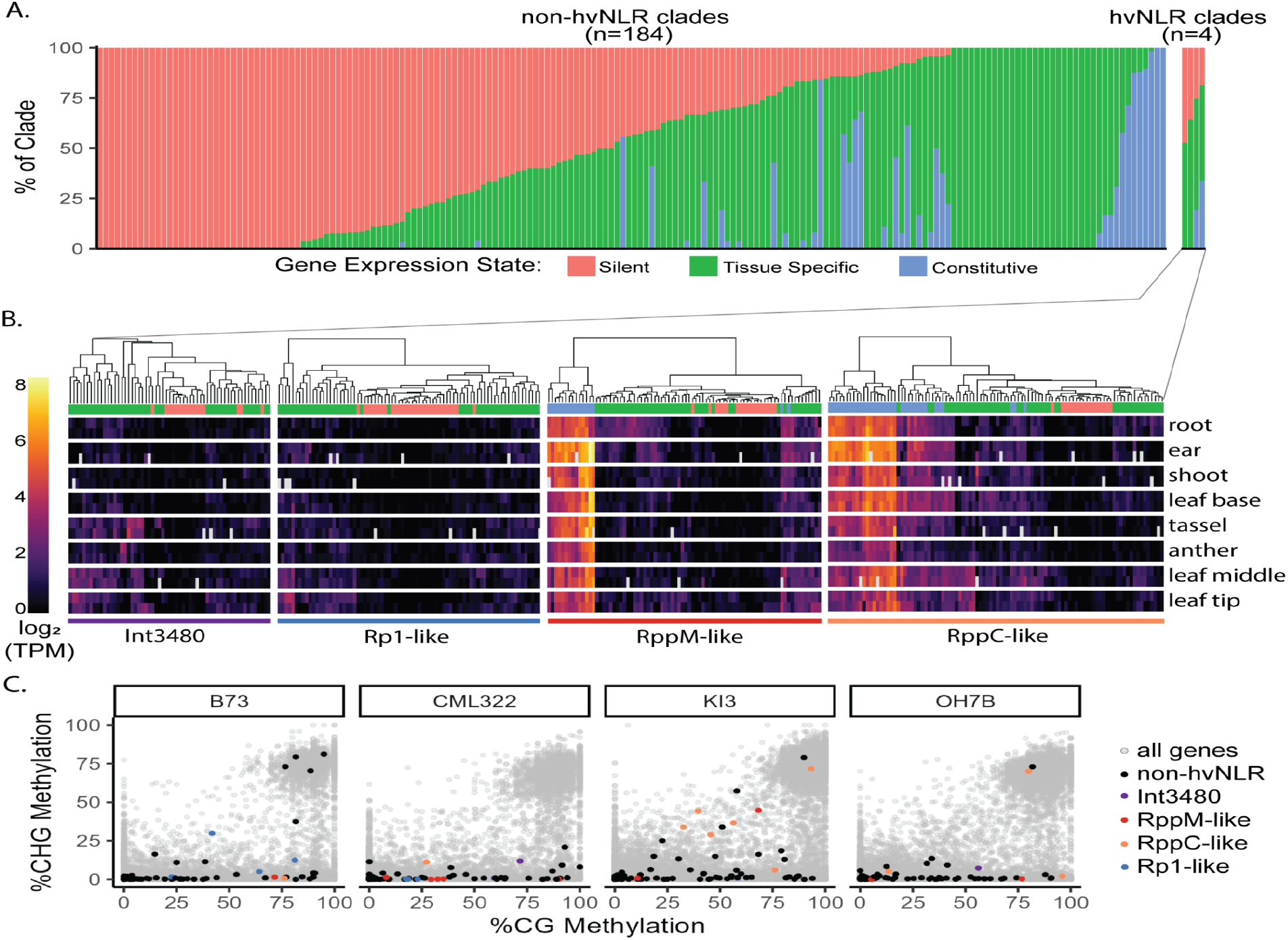
NLRs show variable expression and methylation across lines. **A**. Percentage of each NLR clade that is silent, expressed in a subset of tissues, or constitutively expressed across the 26 NAM lines. **B**. Heatmap of log_2_(TPM) values of hvNLRs, from 8 tissue types with two biological replicates per tissue type shown. Unavailable data is shown in gray. Dendrograms show hierarchical clustering of expression within clades. Expression state of each gene shown as column annotation. **C**. %CG methylation vs %CHG methylation for all genes across four representative NAM lines, with NLRs highlighted.

RppC and RppM confer resistance to southern corn rust (*Puccinia polysora*), an obligate biotroph that primarily infects the upper surface of the leaf (Deng et al. 2022; Wang et al. 2022; Sun et al. 2021). We observe 34% of RppC-like NLRs to be constitutively expressed, 47% to be tissue-specific, and 19% to be silent (Figure 3A). Tissue specific RppC-like NLRs are primarily expressed in the middle and base of the leaf, consistent with the leaf pustule symptoms of southern corn rust (Figure 3B; median log_2_(TPM)=1.64, 1.26 respectively). It has been previously reported that RppM is constitutively expressed in all developmental stages and all tissues in the resistant elite maize inbred line Jing2416K, with the strongest expression in leaves (Wang et al. 2022). Jing2416K is not a NAM founder line, but we observe 19% of RppM-like clade members to be constitutively expressed, 55% to be tissue specific, and 25% to be silent (Figure 3A). Inconsistent with the infection strategy of southern corn rust, the tissue specific RppM-like NLRs are most expressed in root tissue, followed by ear and leaf tip tissue (Figure 3B, median log_2_(TPM) = 1.56, 1.25, and 1.21 respectively). Rp1 confers resistance to common rust, *Puccinia sorghi*, an obligate parasite that primarily infects leaves (Collins et al. 1999). 53% of the Rp1-like clade show tissue specific expression, most often to the middle of the leaf (Figure 3B, median log2(TPM)=1.04), consistent with the infection strategy of *Puccinia sorghi*. 64% of hvNLR clade Int3480 with unknown pathogen specificity is tissue specific, primarily to tassel, middle, and tip of the leaf (Figure 3B, median log_2_(TPM= 1.23, 1.09, and 1.09, respectively).

The varied expression dynamics of members of the same clade may suggest variation in gene body methylation across lines, which has a weak but significant association with expression (Zeng et al. 2023). We calculated the percent of methylated cytosines across the exons of NLRs using enzymatic methyl-seq from seedling tissue (Hufford et al. 2021). Methylation exclusively in the CG context is associated with broad expression and sequence conservation, while both CHG and CG methylation is associated with TE silencing, low expression, and poor sequence conservation (Muyle et al. 2022). We observed across-line variation in NLR methylation dynamics (Figure 3C, Supplemental Figure 3). The majority of NLRs have no CHG methylation, but are distributed along the continuum of CG methylation (Zeng et al. 2023). The amount of CHG methylation varies per line, ranging from several NLRs exhibiting TE-like methylation to only a few to none (Figure 3C, Supplemental Figure 3).

In order to interpret the sequence diversity information for the hvNLR clades, we downloaded AlphaFold2 models of the representative genes from the European Bioinformatics Institute repository (https://alphafold.ebi.ac.uk) and mapped the alignment entropy scores onto the predicted protein surfaces. In all four hvNLR clades we observed that the surface of the LRR domains contained a cluster of variable residues (Figure 4). This pattern of sequence variation suggests a direct recognition mechanism similar to that of RPP1, Roq1, and Sr35, the direct-recognition NLRs with structures presently available in the PDB (Zhao et al. 2022; Martin et al. 2020; Ma et al. 2020; Förderer et al. 2022). In the Int3480 clade, there were two such clusters on the opposite side of the LRR horseshoe that both faced the central cavity, suggesting a bipartite binding site in this receptor (Figure 4A). Similar to Arabidopsis hvNLRs, distantly related hvNLRs in maize differed in the predicted location of candidate ligand binding residues.

**Figure 4.**
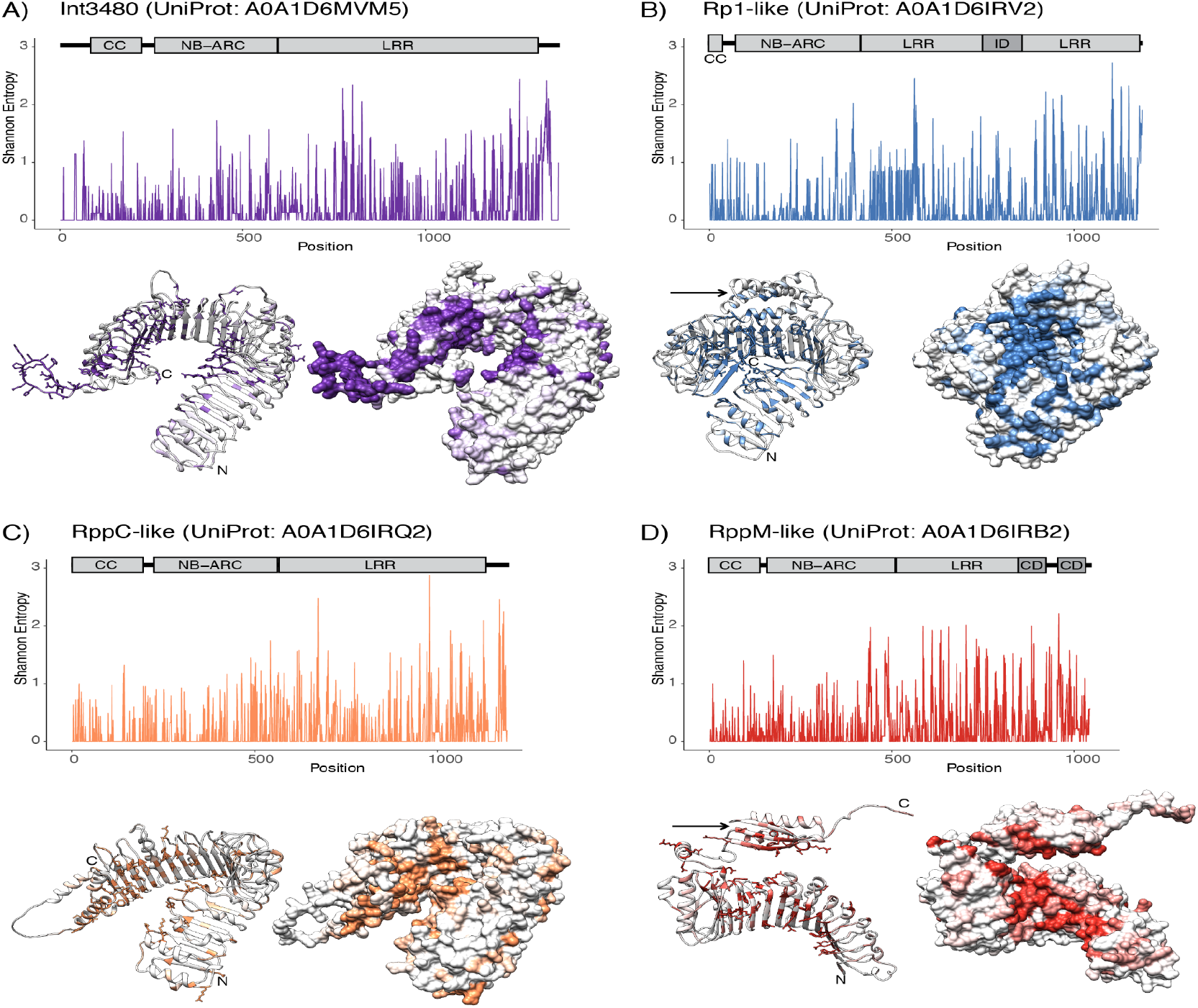
Alignment entropy and AlphaFold2-generated models identify maize hvNLR target-binding sites. In all four clades, the majority of variable residues were mapped to the Leucine-Rich Repeat (LRR) domain. Maize hvNLRs contain additional non-LRR regions with high sequence variability: Rp-1 insertion domain (ID) and RppM capping domain (CD) are highlighted with dark gray boxes on domain diagrams and indicated with arrows on ribbon diagrams. In both ribbon and surface diagrams, the Coiled Coil (CC) and NB-ARC domains are omitted for clarity. Start and end of the post-NB-ARC region are marked with N and C, respectively.

In the Rp1-like clade, we observed an insertion domain nested within the LRR sequence (Figure 4B). The insertion domain is a four-helix bundle with the variable residues clustering on a single face. The non-LRR nature of this sequence was noted by Collins et al (Collins et al. 1999), who wrote that the C-terminal portion of the Rp1-D protein “contains two sections with irregular LRRs separated by a region that cannot be easily arranged into repeats”. Since such an unusual gene structure of Rp-D was determined by direct sequencing of cDNA clones (Collins et al. 1999), the insertion domain has experimental support and is unlikely to be an annotation error. By constructing a Hidden Markov Model (HMM) for this insertion domain and searching against reference proteomes, we found that it is broadly present in Poaceae and is commonly associated with the NB-ARC domain. It therefore appears specific to a clade of NLRs in grasses. By searching for this domain in the AlphaFold2 database, we found maize NLRs and rice NLRs with this ID present as an insertion in the middle of the LRR. Within the NAM lines, Rp1 insertion domain was limited to the Rp1-like clade (Supplemental Figure 4), where it was present in both hvNLR and non-hvNLR subclades.

In RppM-like receptors we found a typical predicted binding site in the LRR region supplemented with what appeared as an additional domain at the LRR C-terminus (Figure 4D). Upon further inspection, this domain turned out to be closely related in sequence to the last two repeats of the contiguous LRR region, which act as a cap on the LRR domain. We investigated a potential evolutionary origin of this domain arrangement and found that sequences with the RppM cap domains are found in 11 neighboring clades, in addition to the RppM-like clade (Supplemental Figure 4). We also observed that the majority of sequences in RppM-like, and in non-hvNLR clades Int4674, Int4977, and Int4916 have two copies of the cap domain. Based on how common this arrangement is in RppM-like and its neighbor clades, we concluded that it is probably not caused by a protein annotation error. Instead, it is likely that incorporating a second copy of the cap allowed receptors in these subfamilies to gain a flexibly attached small domain with a variable surface that can provide additional area for target binding.

In the other two hvNLR clades, the variable residues found outside of LRR included regions that were likely flexible (Figure 4A) or formed alpha-helices that were not part of any predicted domains (Figure 4C). We believe that the overall effect of these non-LRR highly variable sequences might be to increase the overall receptor area available for target binding. Experimental validation will be required to establish any functional role for these non-LRR variable residues and for the Rp1 and RppM insertion domains.

To make analysis of maize NLR sequence diversity broadly accessible, we created a Google Colaboratory notebook that we called NLRCladeFinder. The Colab is available in the https://github.com/daniilprigozhin/NLRCladeFinder repository and includes access to both maize and our published Arabidopsis NLR dataset (Prigozhin and Krasileva 2021). The notebook allows users to input their NLR sequence, find out whether it belongs to a highly variable clade, and to generate an updated clade alignment, phylogenetic tree, and a Chimera annotation file that can be mapped onto a structure model. This interface can be used in tandem with ColabFold (Mirdita et al. 2022), a user-friendly implementation of AlphaFold 2 (Jumper et al. 2021), to obtain structural models and sequence diversity information for maize NLRs.

To illustrate this application, we predicted the structure of the recently cloned Southern corn rust resistance gene RppC (Deng et al. 2022) in ColabFold (Figure 5A) and ran it through NLRCladeFinder (Figure 5B and 5C). Due to limitations in available computational resources, we modeled RppC protein structure from two overlapping fragments. We found that the exact sequence of RppC as published by Deng et al is not present in the NAM pan-proteome. From the updated clade tree, we found that ZM00037AB409540 isoform 2 from the NC358 line is its closest homolog; it differs from RppC by a single R253T substitution. We then combined the predicted model of RppC with a sequence diversity attribute file in Chimera to identify the likely RppC binding site (Figure 5D). This showed a number of exposed hydrophobic residues surrounded by charged residues in the very C-terminal repeats of a large LRR array - features consistent with a protein-protein binding site.

**Figure 5.**
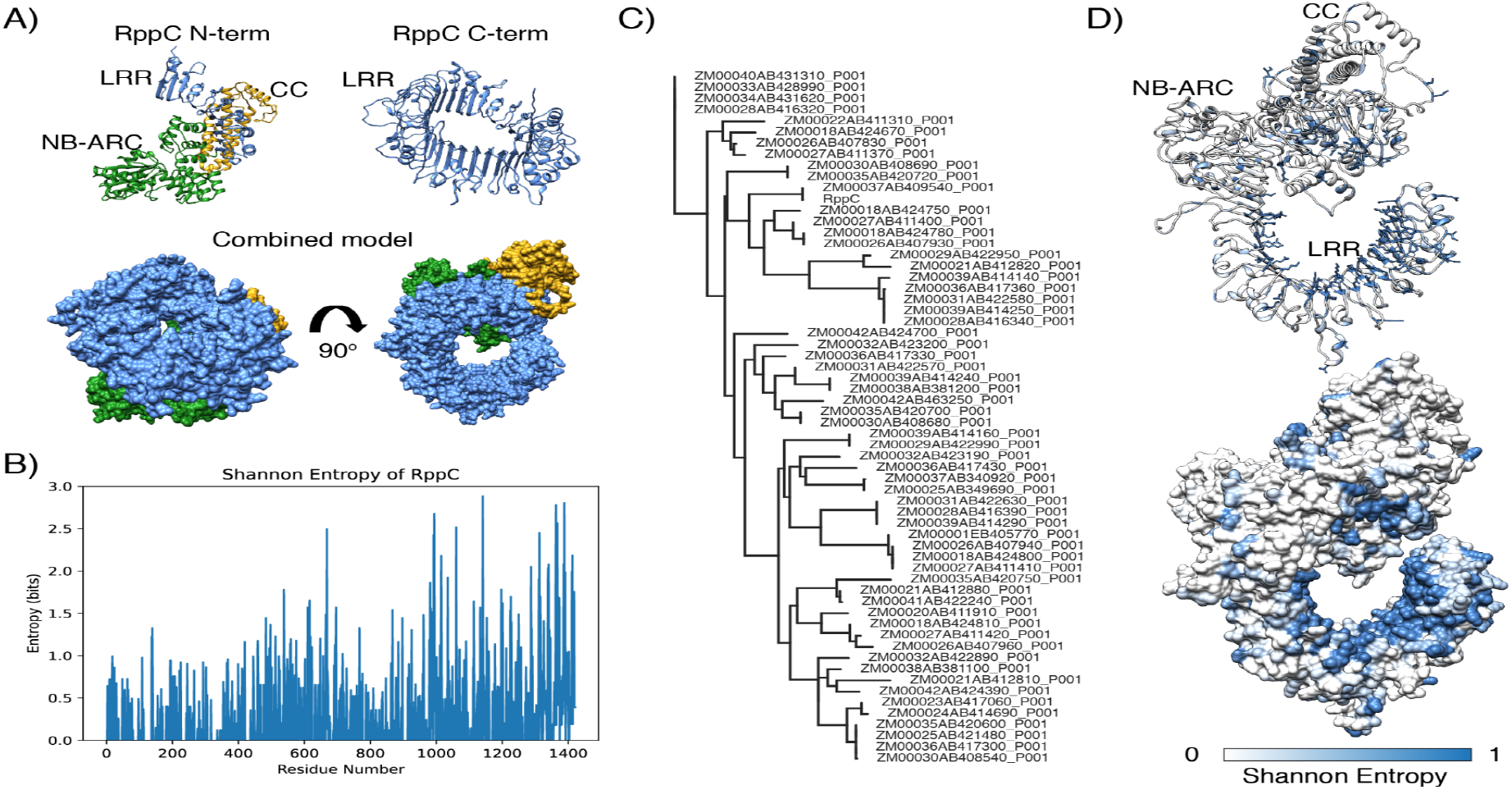
Using NLRCladeFinder and Colab Fold to analyze RppC overall structure and its target-binding site. A) ColabFold-generated model of RppC. Due to limitations on sequence size, we generated two predicted structures spanning residues 1-800 and 620-1422 and combined them in UCSF Chimera (Pettersen et al. 2004). B) Alignment entropy plot generated by NLRCladeFinder. C) NLRCladeFinder-generated tree for the clade containing RppC-like sequences. D) AlphaFold 2 RppC model colored by alignment entropy highlights the likely target binding site in RppC.

## DISCUSSION

Plant immune receptors display a lot of variation compared to housekeeping genes. Presence-absence variation, copy number variation, point mutations, and short insertion/deletion polymorphisms combine to produce a diverse set of receptor specificities in a birth and death process (Michelmore and Meyers 1998). NLR subfamilies differ in the amount of within-species diversity they exhibit (Rose et al. 2004; Bakker et al. 2006; Van de Weyer et al. 2019). Recent advances in long read and enrichment sequencing have enabled family-wide analysis of NLR sequence diversity (Van de Weyer et al. 2019; Seong et al. 2020; Stam et al. 2019). We have previously leveraged these data to highlight the phylogenetic distribution of hvNLRs in *Arabidopsis* and *Brachypodium*, two model species (Prigozhin and Krasileva 2021). Here we applied the same approach to study the NLR immune receptors of maize, a major crop species.

Maize has undergone an overall reduction in the number of NLRs (Hufford et al. 2021). We have found three of its four hvNLR subfamilies, containing Rp1, RppM, and RppC, are within a 2-megabase segment of chromosome 10. This suggests that at least the NAM lines are limited in their NLR diversity compared to *Arabidopsis* and *Brachypodium*, and might thus be a poor source of new NLR-based disease resistance. With growing availability of genomic resources, future work utilizing pan-genome resources for teosinte and other related Andropogoneae species will help determine if this reduction occurred during the domestication process. Given that hvNLRs can present barriers to hybridization (Chae et al. 2014), their relative lack in maize might also contribute to the success of maize hybrids.

NLRs are variably expressed within clades, with close homologs showing constitutive, tissue specific, and silent expression patterns across NAM lines. Regulatory evolution and coding sequence evolution are associated, and each of the four hvNLR clades show a mix of gene expression states and a range of methylation across NAM founder lines, consistent with their rapid amino acid evolution (Davidson et al. 2012; Zeng et al. 2023). This pan-genome, across-tissue NLR expression analysis allows us to investigate recent proposed hypotheses about the relationship between NLR expression and function in maize. High steady state expression is proposed to be signature of functional NLRs (Brabham et al. 2023), and consistent with that theory, a subset of the recently cloned RppM-like and RppC-like NLRs are constitutively expressed and among the most expressed NLRs in leaf tissue across NAM founder lines. Additionally, tissue specific NLR expression may predict pathogen specificity (Contreras et al. 2023; Lüdke et al. 2023). In the NAM founder lines, RppC and Rp1-like clades show tissue specific expression consistent with infection type, but RppM did not. Susceptibility and resistance information for each NAM founder allele is required to validate either of these hypotheses, but the signature of constitutive expression could prioritize alleles for resistance screening, and tissue specific expression could generate hypotheses for the mechanism and pathogen specificity of yet to be described NLR clades such as Int3480.

Recent advances in computational structure prediction have accelerated our understanding of the plant-pathogen evolutionary contest, highlighting both diversity and hidden structural similarities among pathogen effectors (Seong and Krasileva 2021, 2023). Here we used the predicted protein structures of maize NLRs to locate the variable residues and identify these proteins’ likely target-binding sites. These specificity-determining residues characteristically cluster on the concave side of the LRR domain. Despite the differences in exact positions of the candidate binding residues among hvNLRs, our results agree with the inference by Förderer et al that target binding on the inside of the LRR favors an open NB-ARC conformation in an activation mechanism shared among distantly related direct-recognition NLRs. Targeted modification of the highly variable LRR residues might lead to derivation of novel immune specificities in the native maize NLRs, as demonstrated for wheat disease resistance genes Sr33, Sr50 (Tamborski et al. 2023), and Sr35 (Förderer et al. 2022) thus overcoming the relative lack of natural diversity or enable rapid transfer of functional specificities into modern varieties through genome editing of key amino acid sites.

## METHODS

### Pipeline management

To streamline application of HMM sequence searches, phylogeny building, and refinement that underpin our approach to NLR sequence analysis, we implemented our published analysis pipeline (Prigozhin and Krasileva 2021) in the Snakemake workflow management system (Köster and Rahmann 2012). This integrates calls to widely used software including RAxML 8.2.12 (Stamatakis 2014), HMMER 3.3.2 (Eddy 2011), and MAFFT 6.903b (Nakamura et al. 2018) with calls to R scripts (RStudio version 1.4.1106, R version 3.6.0) available at the project github (https://github.com/krasileva-group/maize_nlrome). For all steps, input parameters are provided from and recorded in the Snakemake Snakefile. R packages used for NLR sequence analysis included: entropy (Hausser and Strimmer 2008), tidyverse (Wickham et al. 2019), msa (Bodenhofer et al. 2015), tidytree, ggtree (Yu 2020), treeio, parallel, phangorn, Biostrings, and gggenomes.

### NLR identification and overall phylogeny construction

Maize NLRs were identified using the hmmsearch command from HMMER package version 3.3.2 using a previously described extended NB-ARC domain model (Bailey et al. 2018). Initial alignments of the NB-ARC domain were generated using the –A option, and the NB-ARC alignments of NLRs from different NAM lines were merged into a single fasta-formatted alignment using Easel tools (part of HMMER). The alignment was filtered for sequence length (>30 amino acids) and isoform (isoform 1 of every gene), and a maximum likelihood phylogeny was constructed in RAxML version 8.2.12 (Stamatakis 2014) using PROTCATJTT model with 100 rapid bootstrap replicates (raxml -T 8 -f a -x 12345 -p 12345 -# 100 -m PROTCATJTT). This and the following phylogenetic trees generated in this project can be viewed in iToL (https://itol.embl.de/shared/daniilprigozhin).

### Initial clade identification and sub clade refinement

Scripts - reduce_pfam.R and DomainDiagrams_sm.R - were used to decorate the phylogenetic tree in iToL (Letunic and Bork 2019). Scripts - Zm_NLRome_InitialAssignment_sm.R and Zm_NLRome_PrintInitClades.R - were used to identify a set of initial clades using clade size and clade bootstrap support as key input parameters. Initial clades were then refined by constructing full-length *de novo* protein sequence alignments using MAFFT version 6.903b, phylogeny generation with RAxML version 8.2.12, and selection of well supported nodes for further splitting into sub clades. Candidate node selection was performed using Zm_NLRome_Refinement_sm.R script with branch length, bootstrap support, and NAM line overlap as key parameters. Script Zm_NLRome_PrintRefinedClades.R was then used to convert a set of manually curated split nodes to lists of subclade members for the next round of clade refinement. Refinement converged after 3 iterations.

### Analysis of refined clades and access to clade information through Google Colaboratory

The refined subclades were then analyzed in R using Zm_NLRome_CladeAnalysis.R script, and Shannon entropy values of clade alignment were calculated to identify hvNLR clades with 10 or more positions displaying 1.5 bit or higher entropy. For plotting an individual sequence’s entropy scores, the alignment was filtered to only include columns that have non-gap characters in the reference sequence. In order to make our clade assignments more accessible, we implemented NLRCladeFinder (https://github.com/daniilprigozhin/NLRCladeFinder), a pipeline in Google Collaboratory (Bisong 2019), that can find the best matching clade for a given maize NLR sequence using hmmsearch against a collection of clade-specific HMMs. After a matching clade is found, we use MAFFT (Nakamura et al. 2018) to add the new sequence to the alignment followed by epa-ng (Barbera et al. 2019) and gappa (Czech et al. 2020) to find a phylogenetic placement for the new protein in precomputed clade phylogenies. We also produce a Chimera (Pettersen et al. 2004) annotation file that will let the users display entropy values overlapped on their protein model. The github repository also contains a video tutorial.

### Analysis of NLR Expression

RNA-seq reads from 26 NAM lines and 8 tissue types were downloaded from European Nucleotide Archive (ENA) E-MTAB-8633 and E-MTAB-8628. Reads were mapped to their respective reference genomes using STARv2.7.10a in quant mode with default parameters (Dobin et al. 2013). Counts were converted to transcripts per million (TPM) and averaged across four biological replicates, then log_2_(TPM+1) transformed for visualization. NLRs are repetitive and often similar, making them difficult to sequence with short reads. To determine if any NLRs were unmappable, RNAseq reads were simulated using Polyester v1.2.0 (Frazee et al. 2015). 12 NLRs were subsequently excluded from analysis: ‘Zm00039ab351270’, ‘Zm00026ab135540’, ‘Zm00036ab418650’, ‘Zm00001eb091500’, ‘Zm00001eb164880’, ‘Zm00001eb343890’, ‘Zm00001eb391100’, ‘Zm00001eb405870’, ‘Zm00001eb405900’, ‘Zm00001eb405930’, ‘Zm00033ab429000’, ‘Zm00029ab367660’). To define the gene expression categories, we compared TPM values across the 8 tissues (leaf tip, leaf middle, leaf base, root, shoot, ear, anther, tassel). We used previously defined categories of tissue-specific expression as TPM ≥ 1 in at least 1 tissue and TPM < 1 in at least 1 tissue, constitutive expression as TPM ≥ 1 in all 8 tissues, and silent as TPM < 1 in all 8 tissues (Zeng et al. 2023).

### Analysis of NLR methylation

Enzymatic methyl-seq (EMseq) reads from the second leaves of 26 NAM lines were downloaded from ENA E-MTAB-10088. Reads were trimmed using Trim Galore! V0.6.6 with a Phred cutoff score of 20 and Illumina adapter sequences, with a maximum trimming error rate of 0.1 (Babraham Bioinformatics). Bismarck v0.23.0 was used to prepare each NAM genome, map the EMseq reads, remove duplicates, and determine percent methylation at each cytosine (Krueger and Andrews 2011). Cytosines with at least 5 reads were used for analysis, and symmetrical cytosines within CG base pairs were averaged. The percent methylation of each cytosine site was averaged across the exons of each gene to exclude potential transposable element associated methylation potentially present in the intron sequences (Zeng et al. 2023) and weighted by the number of covered cytosine sites.

## Data and code availability

Data including phylogenetic trees, alignments, expression and methylation values, and files used to set up the snakemake pipeline for this project are archived in Zenodo (DOI: 10.5281/zenodo.11069065). Scripts used in the project are deposited in GitHub (https://github.com/krasileva-group/maize_nlrome). NLRCladeFinder can be found at https://github.com/daniilprigozhin/NLRCladeFinder. Phylogenetic trees generated during this project can also be viewed in iToL (https://itol.embl.de/shared/daniilprigozhin).

## ACKNOWLEDGEMENTS

We are grateful to Dr. Erin L. Baggs, Kyungyong Seong, China Lunde Shaw, and Dr. Wei Wei for the critical reading of the manuscript. We thank Dr. Marc Allaire and members of the Berkeley Center for Structural Biology for their support and the use of computational resources. The Berkeley Center for Structural Biology is supported by the Howard Hughes Medical Institute, the National Institutes of Health, and through participating research team partnerships. This research used the Savio computational cluster resource provided by the Berkeley Research Computing program at the University of California, Berkeley (supported by the UC Berkeley Chancellor, Vice Chancellor for Research, and Chief Information Officer). Ksenia V Krasileva and the work in Krasileva lab has been funded by NIH Director’s Award (1DP2AT011967-01), Gordon and Betty Moore Inventor Fellowship (grant number: 8802) and the Innovative Genomics Institute.

---

**Supplemental Figure 1.**
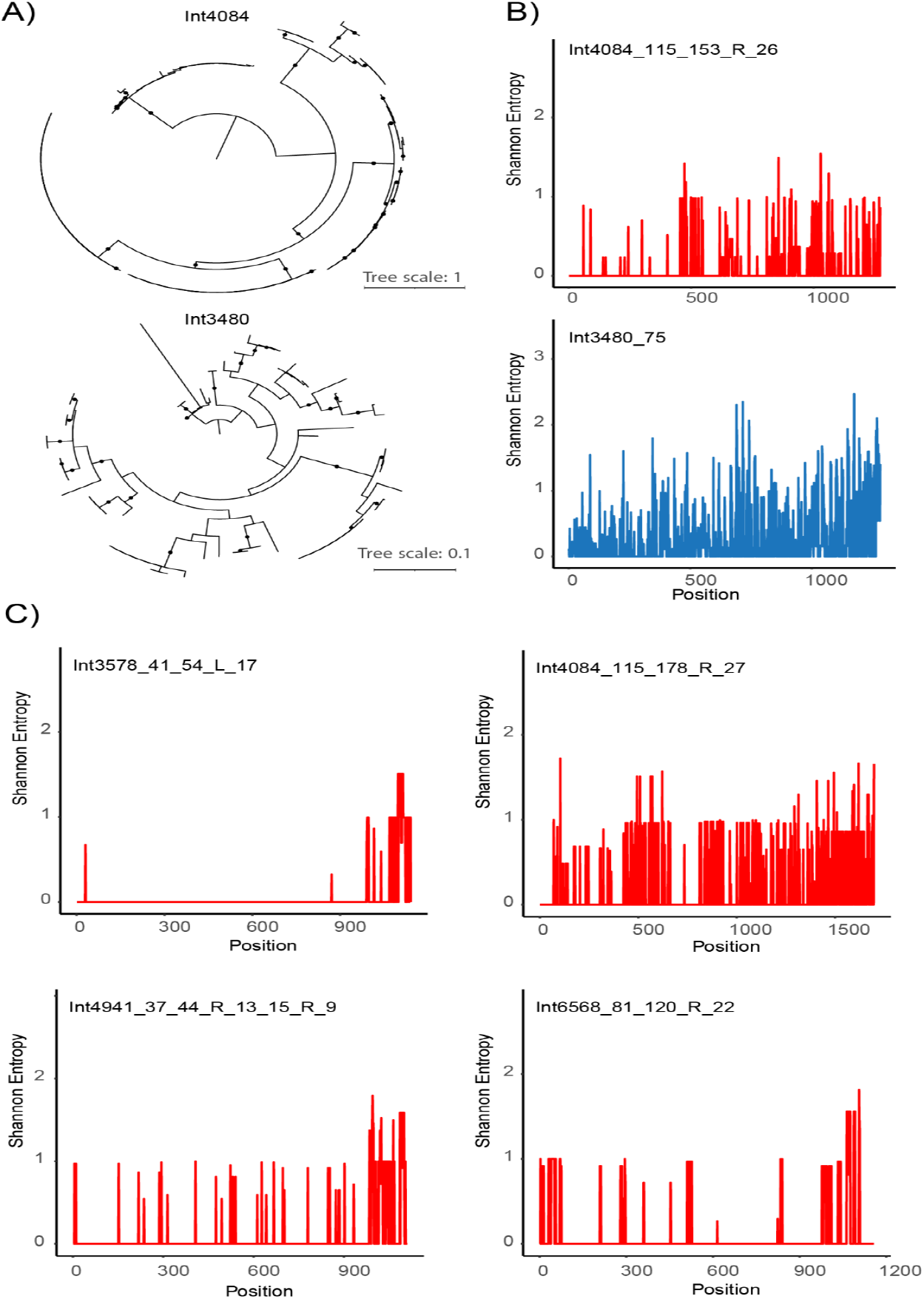
Examples of NLR clade refinement A) Example of two clades from the NLR phylogeny, where Int4084 can be further refined to yield five low-entropy subclades and Int3480 cannot be further split and yields a single high-entropy clade (notice the difference in tree scale). B) Shannon entropy of sequence alignments of two refined subclades showing refined non-hvNLR clade (red) and hvNLR subclade (blue). C) Entropy plots of alignments of NLR clades that satisfy the hvNLR threshold established on Arabidopsis data, but were excluded from the hvNLR analysis. Excluding these clades can also be accomplished by raising the cut off to 20 amino acid sites over the 1.5 bit entropy cutoff.

**Supplemental Figure 2:**
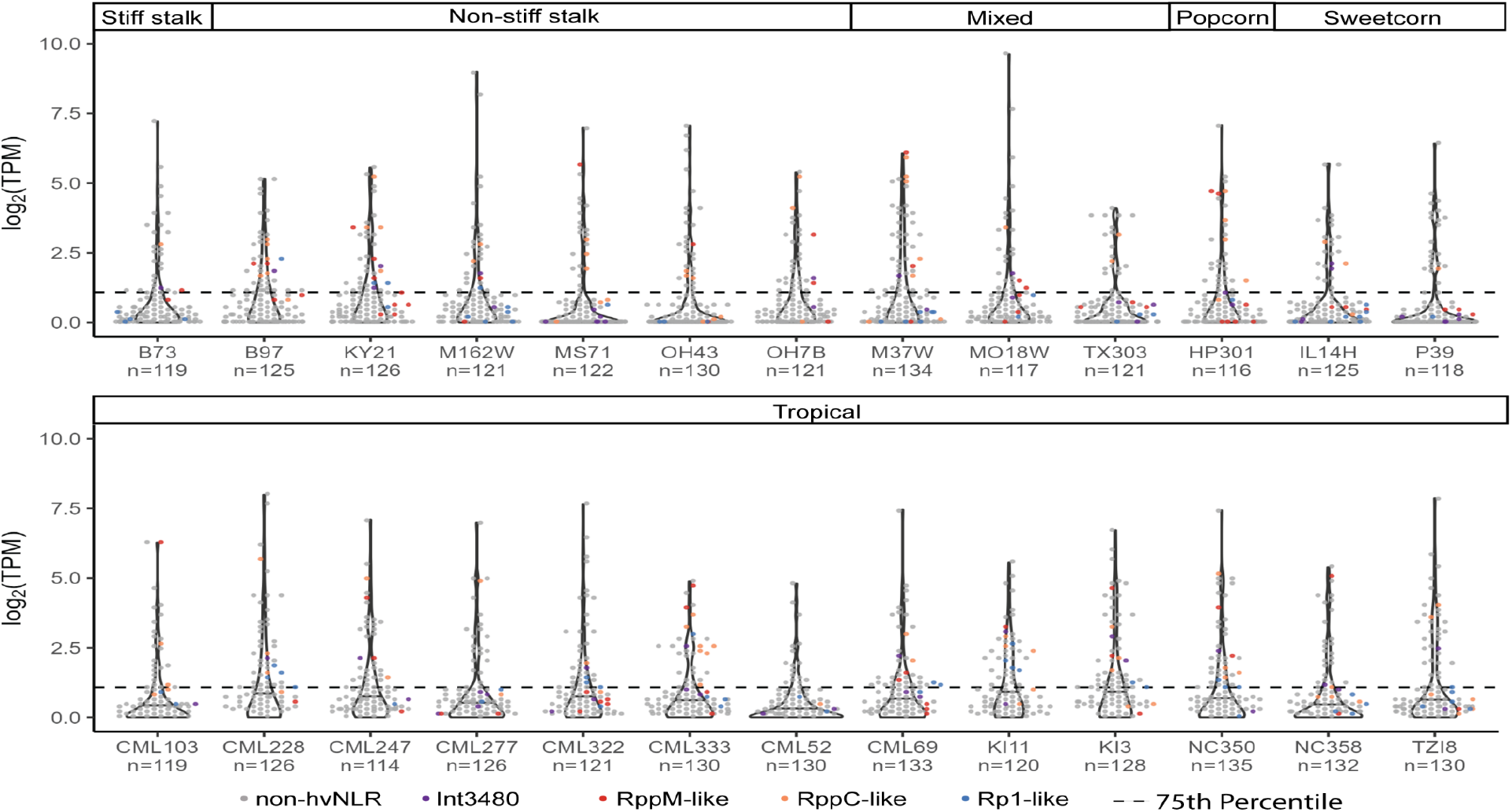
Per-line NLR expression in leaf tissue sampled from the middle of the tenth leaf. NAM lines are grouped by subpopulation membership. hvNLR clades are highlighted. The dashed line represents the 75th percentile of all NLR expression in the middle of the tenth leaf.

**Supplemental Figure 3:**
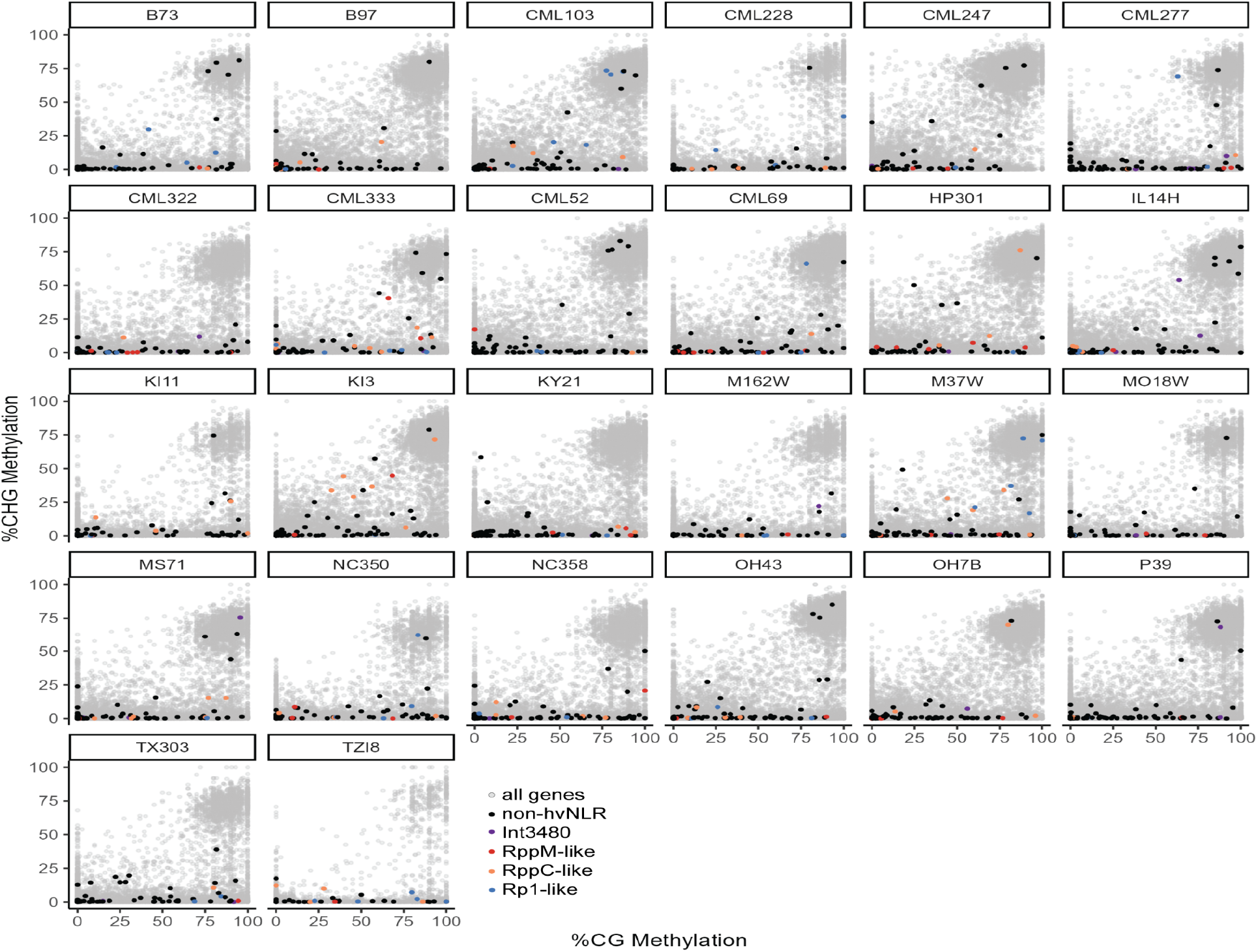
%CG methylation vs %CHG methylation for all genes of all NAM lines, with NLRs highlighted.

**Supplemental Figure 4.**
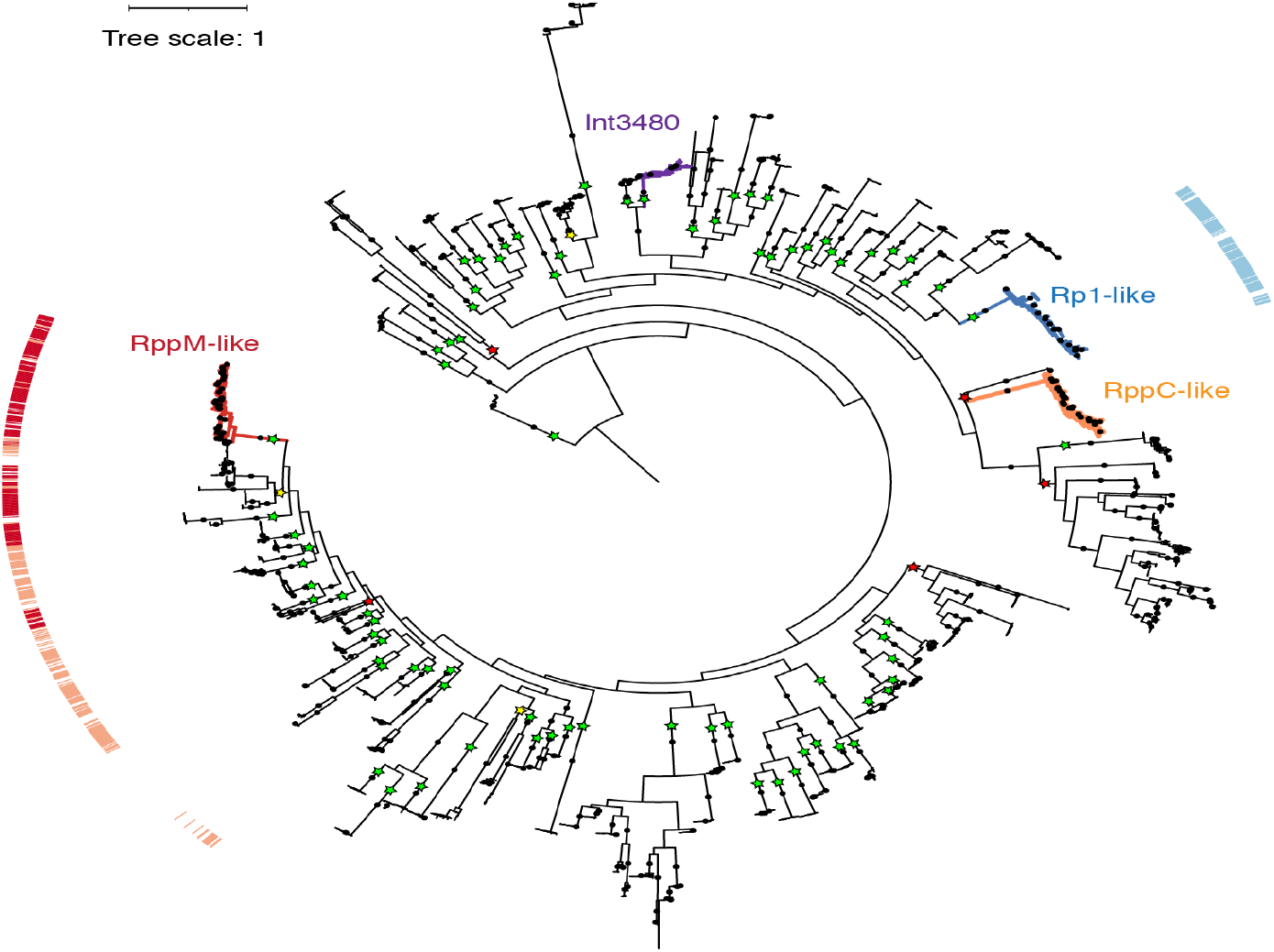
Tree adapted from Figure 1B shows that Rp1 insertion domain (blue ribbon) is limited to Rp1-like initial clade, while RppM capping domain can be present in two copies (red ribbon) or one copy (orange ribbon) and is widespread outside of the RppM-like initial clade.

